# Attentionally modulated subjective images reconstructed from brain activity

**DOI:** 10.1101/2020.12.27.424510

**Authors:** Tomoyasu Horikawa, Yukiyasu Kamitani

**Affiliations:** Department of Neuroinformatics, ATR Computational Neuroscience Laboratories, Kyoto, Japan; Graduate school of Informatics, Kyoto University, Kyoto, Japan

**Keywords:** vision, attention, decoding, visual image reconstruction, fMRI, deep neural network

## Abstract

Visual image reconstruction from brain activity produces images whose features are consistent with the neural representations in the visual cortex given arbitrary visual instances [1–3], presumably reflecting the person’s visual experience. Previous reconstruction studies have been concerned either with how stimulus images are faithfully reconstructed or with whether mentally imagined contents can be reconstructed in the absence of external stimuli. However, many lines of vision research have demonstrated that even stimulus perception is shaped both by stimulus-induced processes and top-down processes. In particular, attention (or the lack of it) is known to profoundly affect visual experience [4–8] and brain activity [9–21]. Here, to investigate how top-down attention impacts the neural representation of visual images and the reconstructions, we use a state-of-the-art method (deep image reconstruction [3]) to reconstruct visual images from fMRI activity measured while subjects attend to one of two images superimposed with equally weighted contrasts. Deep image reconstruction exploits the hierarchical correspondence between the brain and a deep neural network (DNN) to translate (decode) brain activity into DNN features of multiple layers, and then create images that are consistent with the decoded DNN features [3, 22, 23]. Using the deep image reconstruction model trained on fMRI responses to single natural images, we decode brain activity during the attention trials. Behavioral evaluations show that the reconstructions resemble the attended rather than the unattended images. The reconstructions can be modeled by superimposed images with contrasts biased to the attended one, which are comparable to the appearance of the stimuli under attention measured in a separate session. Attentional modulations are found in a broad range of hierarchical visual representations and mirror the brain–DNN correspondence. Our results demonstrate that top-down attention counters stimulus-induced responses and modulate neural representations to render reconstructions in accordance with subjective appearance. The reconstructions appear to reflect the content of visual experience and volitional control, opening a new possibility of brain-based communication and creation.

## Results

We investigated how neural representations regulated by top-down attention overrides those of external stimuli by reconstructing visual images from fMRI responses while subjects paid attention to one of two overlapping images. We collected fMRI data from five subjects in two types of experimental sessions. In the training session, in which the data for model training were collected, subjects passively viewed presented natural images while fixating the center of images (6,000 trials). The test session, in which the data for testing the trained models were collected, consisted of single-image trials and attention trials. In the single-image trials, subjects viewed presented images (10 unique images not included in the stimuli of the training session) as in the training session. In the attention trials, subjects viewed superpositions of two different images (all the 45 pairs from the 10 images) and were asked to attend to one of two superimposed images while ignoring the other such that the attended image was perceived more clearly (Figure 1A).

**Figure 1.**
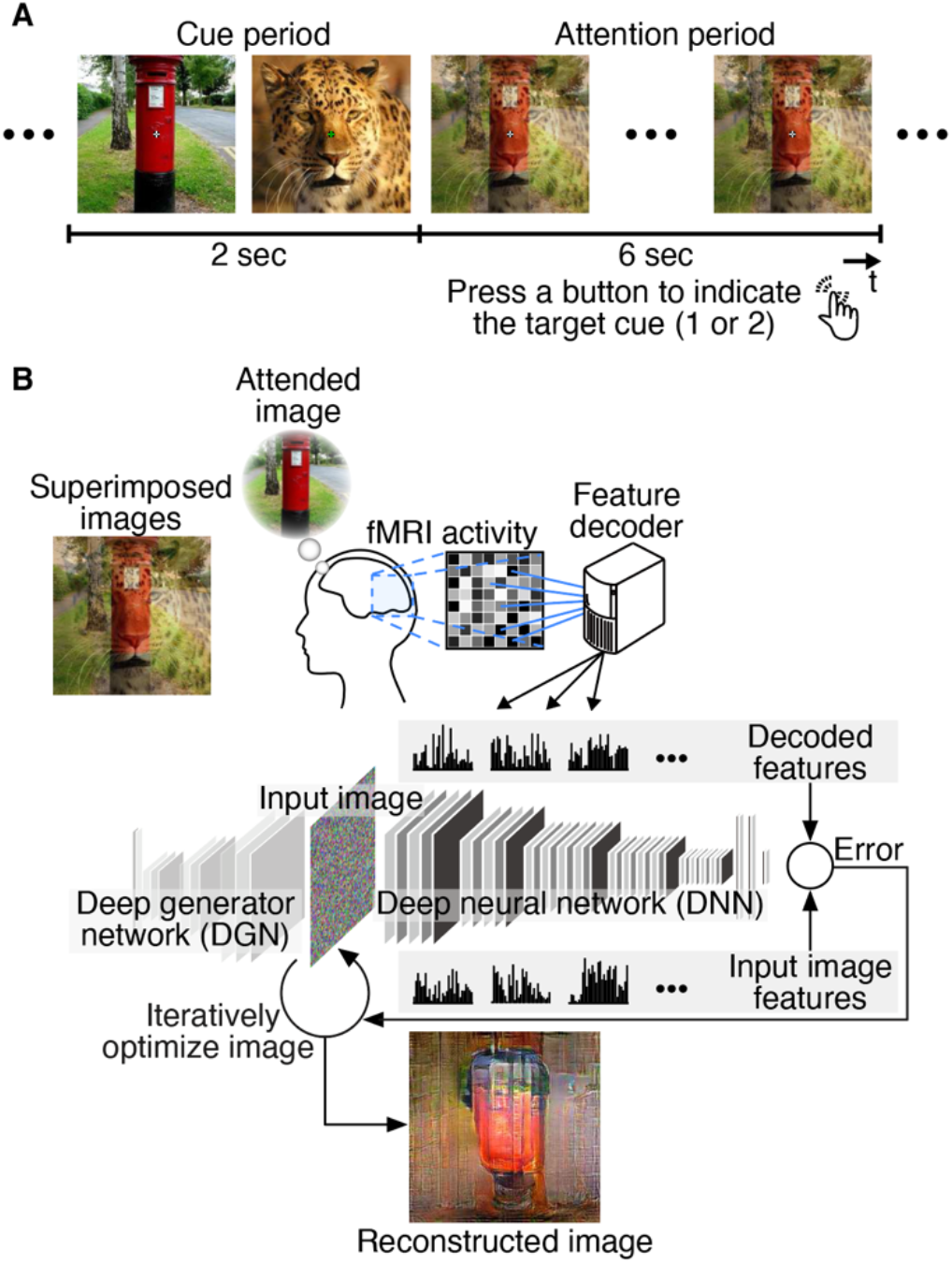
Overview of image reconstruction from brain activity during attention. (A) Experimental design of attention trials. In each trial, two cue images and a superposition of the preceding two cue images (flashed at 2 Hz) were sequentially presented to subjects. During an attention period, subjects were asked to attend to one of two superimposed images indicated by green fixation color during cue periods while ignoring the other. Subjects pressed a button to indicate which of the first or second image were attended to for confirmation. (B) Reconstruction procedure. Given a set of decoded features for all DNN layers as a target of optimization, the method [3] optimizes pixel values of an input image so that the features computed from the input image become closer to the target features. A deep generator network (DGN) [25] was introduced to produce natural-looking images, in which optimization was performed at the input space of the DGN (see Methods: “Visual image reconstruction analysis”).

We analyzed the fMRI data using the deep image reconstruction approach with which we had demonstrated that perceived and imagined images were reliably reconstructed [3]. We first extracted DNN features using the VGG19 model [24] from the images presented in the training session. Then linear regression models (decoders) were trained to predict (decode) the individual DNN feature values from the patterns of fMRI voxel values in the visual cortex (VC) covering from V1 through the ventral object-responsive areas. The trained decoders were then tested on the data from the test session (160 single-image trials and 720 attention trials). The decoded DNN features from each of the single-image and attention trials were processed with the optimization procedure to create a reconstructed image (Figure 1B) [3].

Reconstructions from samples of attention trials are shown in Figure 2A (see Figure S1 for validations of decoders; see Figure S2 and Video S1 for more examples). The generated images appear to resemble the attended images; they tend to represent the shapes, colors, and finer patterns (e.g., faces) of the attended images to a greater degree than those of the unattended images. Notably, even for identical stimulus images, the appearance of reconstruction was strikingly different depending on the attention. The quality of successful reconstructions from attention trials seems comparable to that from single-image trials (Figure 2B).

**Figure 2.**
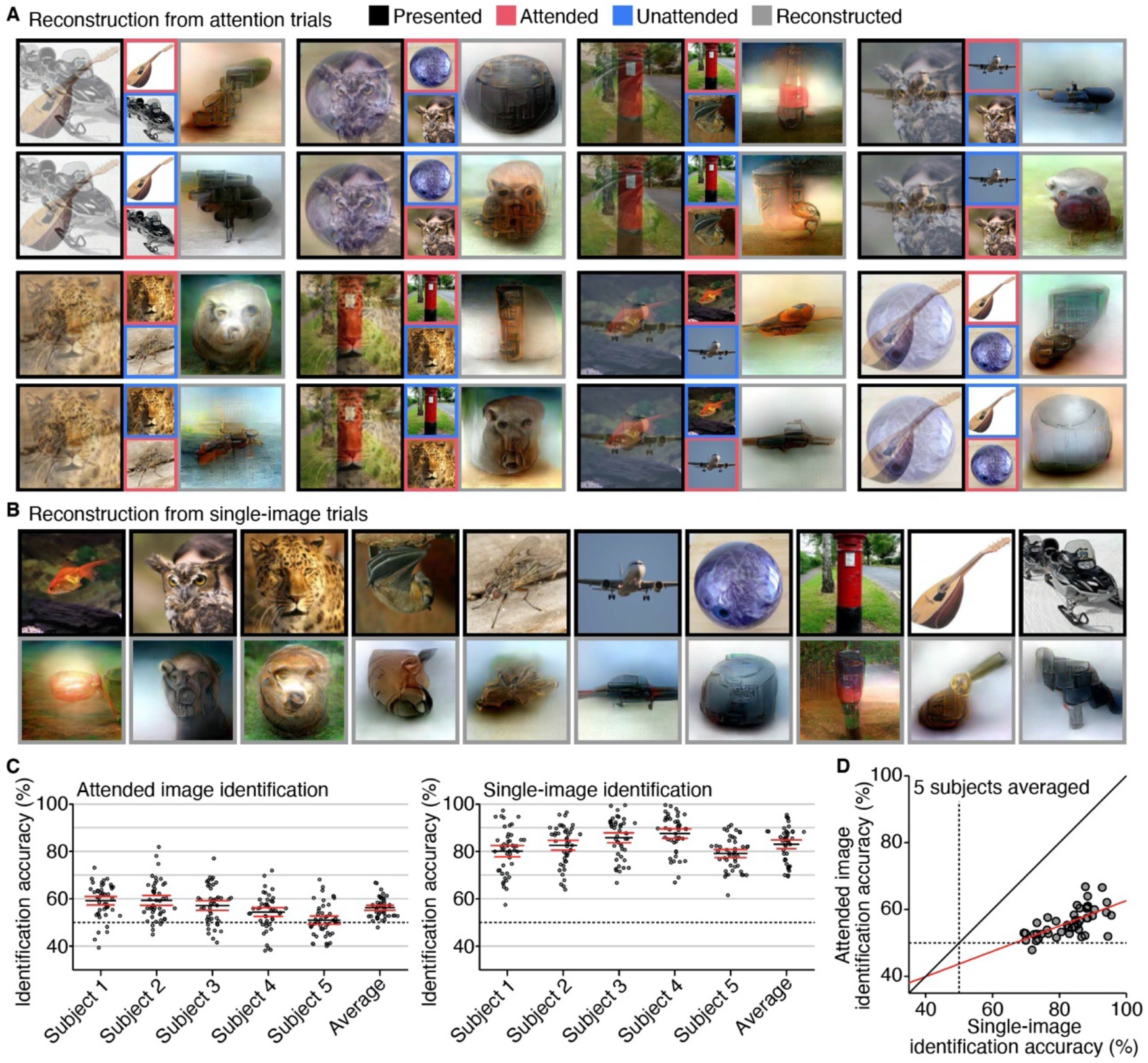
Reconstructions from attention and single-image trials. (A) Reconstructions from attention trials. Reconstructions with relatively high rating accuracies are shown (see Figure S2 for more examples; see Methods: “Evaluation of reconstruction quality”). For each specific presented image, two reconstructions from the same subjects are shown for trials with different attention targets. (B) Reconstructions from single-image trials. Images with black and gray frames indicate presented and reconstructed images, respectively (see Figure S1C for more examples). (C) Identification accuracy based on behavioral evaluations. Dots indicate mean accuracies of pair-wise identification evaluations averaged across samples for each paired comparison (chance level, 50%; see Methods: “Evaluation of reconstruction quality”). Black and red lines indicate mean and lower/upper bounds of 95% C.I. across pairs. (D) Scatter plot of attended and single-image identification accuracy based on behavioral evaluations. Dots indicate mean accuracies averaged over samples for each paired comparison and subjects. The red line indicates the best linear fit.

We evaluated the reconstruction quality by behavioral ratings with a pair-wise identification task. Human raters judged which of two candidates (attended and unattended images for attention trials; true [presented] and false images for single-image trials) is more similar to the reconstruction. Twenty raters performed the identification for each reconstruction with a specific candidate pair (e.g., “post” and “leopard” for a reconstruction with target “post”). The accuracy for each reconstruction can be defined by the correct identification ratio among all raters. However, we will present results using the pooled accuracy for each image pair for attention trials, that is, the correct identification ratio across all raters and the reconstructions (trials) with the same image pair. For each image pair, the accuracies were pooled over the two attention conditions (attend to one or to the other) to cancel the potential effect of image saliency: if identification solely depends on the relative saliency of the two component images regardless of attention, the pooled identification ratios would cancel to the chance level. To be comparable, the accuracy for single-image trials was calculated by pooling the identification results over each pair of single-image stimuli. For both attended image and single-image reconstructions, the mean identification accuracies were significantly higher than the chance (Figure 2C; 56.2%, 95% confidence interval [C.I.] across pairs [55.1, 57.3] for attention; 83%, C.I. [81.1, 84.8] for single-image; five subjects averaged; one-sided Wilcoxon signed-rank test, *p* < 0.01). The accuracy levels for attention trials were modest on average (see Figure S2C and D for failed reconstructions). However, the accuracies were positively correlated between attention and single-image trials across pairs (Figure 2D; *r* = 0.650; *t*-test, *p* < 0.01; correlation from mean accuracies of five subjects). This indicates that attentional modulation was more pronounced for pairs of the images that were easy to decode or reconstruct. Note also that the subjects better at single-image reconstruction did not necessarily achieve high accuracies for attended image reconstructions (e.g., Subject 4). The variances across subjects may reflect individual differences in the capability to exert attention.

Attention is known to enhance the perceived contrast of stimuli [6–8]. We thus sought to model the reconstructed images by superimposed images with biased contrasts (Figure 3A). We created superpositions with weighted contrasts ranging from 0% vs. 100% to 100% vs. 0% (attended vs. unattended) for each pair, where 50% vs. 50% corresponds to the contrasts used for the stimuli in the attention trials. These weighted superpositions were given to the DNN to obtain their stimulus features, which were then compared with neural feature representations. For each DNN layer, Pearson correlations were calculated between the decoded feature pattern from each attention trial and a set of DNN feature patterns of the superimposed images with different contrasts. The weighted contrast that yielded the highest correlation was considered to indicate the degree of attentional modulation. We could use the DNN features calculated from the reconstructions instead of the decoded features, but they highly resembled and yields similar results in this analysis. Thus, the decoded features can be seen as the stimulus features of the reconstructions.

**Figure 3.**
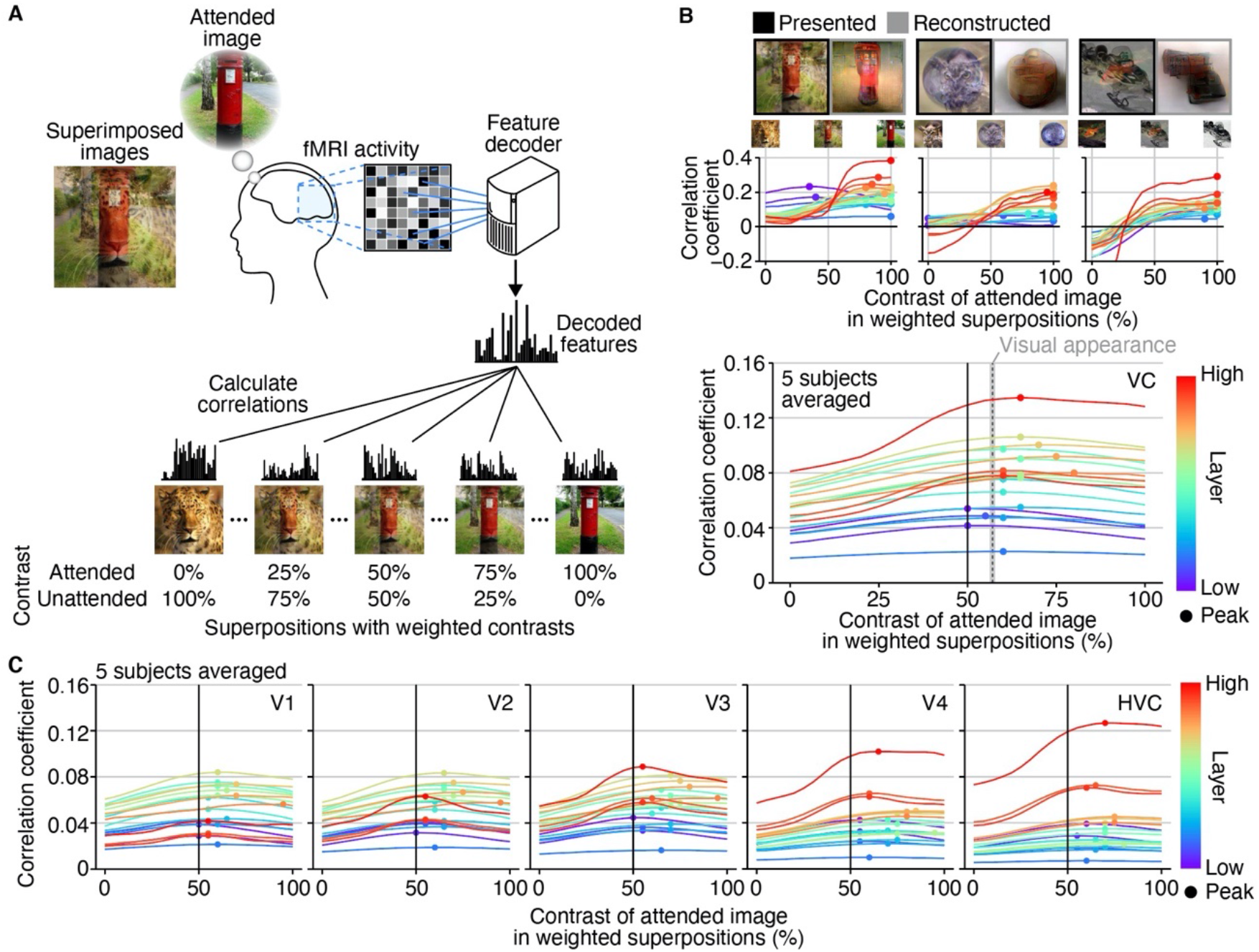
Attentional modulation modeled by image contrast. (A) Evaluation procedure by weighted image contrasts. Correlations were calculated between decoded feature patterns and feature patterns computed from superpositions with weighted contrasts (every 5% steps ranging from 0% vs. 100% to 100% vs. 0% for attended and unattended images; presented images, 50%). (B) Correlation coefficients between decoded feature patterns and feature patterns computed from weighted superpositions. Top panels show results from individual trials with presented and reconstructed images. The bottom panel shows the results averaged across all trials. Colored lines indicate correlations for individual DNN layers (a total of 19 layers; decoded from VC; five subjects averaged; see Figure S3 for the results of individual subjects). Dots indicate contrasts showing the highest correlations with decoded feature patterns. A dashed line indicates the mean contrast of visual appearance evaluated in an independent behavioral experiment (gray area, 95% CI across pairs; see Methods: “Evaluation of visual appearance”). (C) Correlation coefficients between decoded feature patterns from individual visual subareas and feature patterns computed from weighted superpositions. Conventions are the same as in (B).

The decoded features for successful reconstructions of attended images generally showed peak correlations with greater contrasts of the attended images (Figure 3B top; decoded from VC). The estimated correlations often peaked at 100% (i.e., attended image), indicating that representations regulated by top-down voluntary attention can override those from external stimuli. Overall, the peaks of the correlations were shifted toward attended images in most DNN layers except for some lower layers (Figure 3B bottom; 62.6%, mean across layers; averages across trials and five subjects). In an independent behavioral experiment, we measured the perceived contrasts of equally weighted stimuli under attention by matching the stimulus contrasts after the attention period. The matched contrast (indicated by “visual appearance” in Figure 3B bottom) was relatively small but comparable to the biases observed in the decoded features (57.0%, three subjects averaged; see Methods: “Evaluation of visual appearance”).

An additional analysis using five visual subareas (V1–V4 and higher visual cortex [HVC]) showed similar results even with lower visual areas: the peak shift toward the attended image was observed in 403 out of 475 (= 5 subjects × 19 layers × 5 areas) conditions (84.8%; Figure 3C; see Figure S3 for the results of individual subjects). These results indicate that robust attentional modulations are found across visual areas and the levels of hierarchical visual features as measured by the equivalence to biased stimulus contrasts.

Finally, we further investigated the attentional modulations in terms of the feature specificity in individual visual areas. Here, we performed a pair-wise identification analysis based on feature correlation, in which a decoded feature pattern was used to identify an image between two candidates by comparing the correlations to the image features. The identification of attended images was performed for all combinations of areas and layers, and the results were compared with single-image identification (Figure 4A).

**Figure 4.**
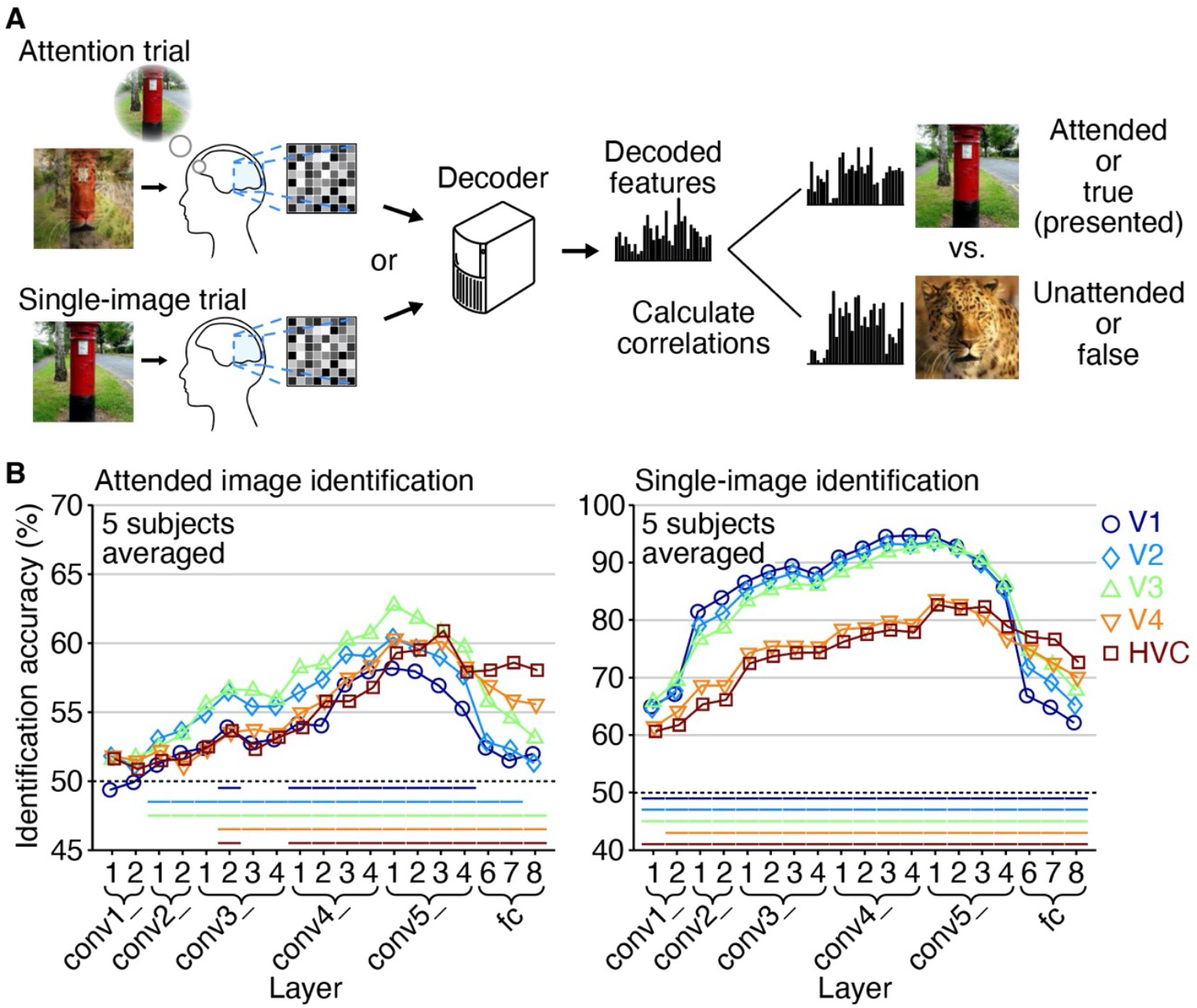
Identification by feature correlation. (A) Identification procedure. Pair-wise identification of attended images and single presented images were performed via decoded features obtained from each sample of the attention and single-image trials (chance level, 50%; see Methods: “Identification analysis”). (B) Identification accuracy from attention and single-image trials. The analysis was performed for all combinations of individual visual subareas and DNN layers (VGG19). Colored lines beneath data indicate results of statistical tests (for attention trials, one-sided binomial test, *p* < 0.05 for three out of five subjects; for single-image trials, one-sided Wilcoxon signed-rank test, *p* < 0.05 for all subjects; see Figures S4 for the results of individual subjects).

While V1–V3 show markedly superior performance in single-image identification especially at lower-to-middle DNN layers, such superiority is diminished in attended image identification. As for V1, the attended image identification is generally poor across all DNN layers. Thus, V1– V3 appear to play a major role in representing stimuli, but not as much in attentional modulation. The attention identification performances of different brain areas show similar profiles peaking across middle-to-higher DNN layers. The representations of these levels may be critical in attentional modulation.

A closer look reveals a hierarchical correspondence between the brain areas and DNN layers. The attended image identification shows higher accuracies from lower-to-middle areas (V2 and V3) with features of lower-to-middle layers (conv2–5) and from higher areas (V4 and HVC) with features of higher layers (fc6–8; Figure 4B left; see Figure S4 for the results of individual subjects). This accuracy pattern generally mirrors the tendency found in the single-image identification performance (Figure 4B right) except V1. These results suggest that attentional modulation is also constrained by the hierarchical correspondence between brain areas and DNN layers for stimulus representation.

## Discussion

In this study, we investigated how top-down attention modulates the neural representation of visual stimuli and their reconstructions using the deep image reconstruction approach. We found that the reconstructions from visual cortical activity during selective attention resembled the attended images rather than the unattended images. While reconstruction quality varied across stimuli and trials, successful reconstructions stably reproduced distinctive features of attended images (e.g., shapes and colors). When the reconstructions were modeled using superimposed images with biased contrasts, attentional biases were observed consistently across the visual cortical areas and the levels of hierarchical visual features and were comparable to the appearance of equally weighted stimuli under attention. The identification analysis based on feature correlations revealed elevated attentional modulation for middle-to-higher DNN layers across the visual cortical areas. Attentional modulation exhibited a hierarchical correspondence between visual areas (except V1) and DNN layers, as found in stimulus representation. Our analyses demonstrate that top-down attention can render reconstruction in accordance with subjective experience by modulating a broad range of hierarchical visual representations.

We have shown robust attention-biased reconstructions especially with image pairs whose individual images were well reconstructed when presented alone (Figure 2D). However, there were substantial performance differences across subjects. We found that subjects with higher performances in single-image reconstructions (e.g., Subject 4) did not necessarily show better attended image reconstructions (Figure 2C) nor stronger attentional modulations (Figures S3 and S4). The difference of attentional modulation across subjects may be attributable to the individual difference in the capability to control attention. Exploring psychological and neuronal covariates with these differences can be an important research direction for future studies.

Reconstructions were explained by superimposed images with contrasts biased to the attended ones, which were comparable to the appearance of stimulus images under attention (Figure 3B). On average, the decoded features were most correlated with the stimulus features with biased contrasts around 60% vs. 40%, overriding the 50% vs. 50% contrasts in the stimuli. However, it should be noted that the peak biases were variable across DNN layers as well as visual areas, image pairs, and trials. Further, the visual features of biased stimulus images cannot account for the interaction of attentional modulations across layers. Thus, biased image contrast should be considered a rough approximation of attentional modulation in the visual system.

Previous studies of attention mainly focused on a few specific aspects/features of interest (e.g., semantic categories [12, 16–19], edge orientations [13], motion directions [14], properties of receptive field models [19, 20]) to investigate the effects of attentional modulations on neural representations. In contrast, our approach is based on hierarchical DNN features that are discovered via the training with a massive dataset of natural images. This allows us to examine attentional effects on hierarchically organized visual features that are difficult to be designed by an experimenter. Furthermore, the image reconstruction from decoded features enables in-depth examinations of the extent and specificity of attentional effects. As this approach primarily relies on the validity of DNNs as computational models of the neural representation [26, 27], the use of more brain-like DNNs [28, 29] may enhance the efficacy to reveal fine-grained contents of attentionally modulated visual experience.

A limitation of this study is the lack of explicit instructions to subjects about the strategy for directing their attention to target images, which might partly explain the variations across subjects (Figure S4C). Higher visual areas tended to be more closely linked to attentional modulation (Figure 3C) potentially because the subjects paid attention more to categorical aspects of the stimulus. Future experiments with explicit instructions regarding attention strategy would elucidate how specifically and flexibly attention can be deployed.

The present study analyzed fMRI activity in a single trial-based manner. Despite the relatively low signal to noise ratio, the reconstructions were of comparable equality to those from trial-averaged data [3]. The single trial-based reconstruction of subjective images opens new possibilities of applications. It can be applied to new experimental designs that use real-time decoding/reconstruction and the feedback of the information. Furthermore, as the reconstruction reflects the content of experience and volitional control, it may provide a new means to express and communicate internal messages in the form of visual images.

## ACKNOWLEDGMENTS

The authors thank Misato Tanaka for assistance with the reconstruction evaluation experiment. This study was conducted using the MRI scanner and related facilities of Kokoro Research Center, Kyoto University. This research was supported by grants from the New Energy and Industrial Technology Development Organization (NEDO), JSPS KAKENHI Grant number JP15H05710, JP15H05920, JP17K12771, and 20H05705. JST CREST Grant Number JPMJCR18A5, and JST PRESTO Grant Number JPMJPR185B Japan.

## AUTHOR CONTRIBUTIONS

Conceptualization, T.H. and Y.K.; Methodology, T.H.; Validation, T.H.; Formal Analysis, T.H.; Investigation, T.H.; Resources, T.H. and Y.K.; Writing – Original Draft, T.H.; Writing – Review & Editing, T.H. and Y.K.; Visualization, T.H.; Funding Acquisition, T.H. and Y.K.

## DECLARATION OF INTERSTS

The authors declare no competing interests.

## METHODS

### KEY RESOURCES TABLE

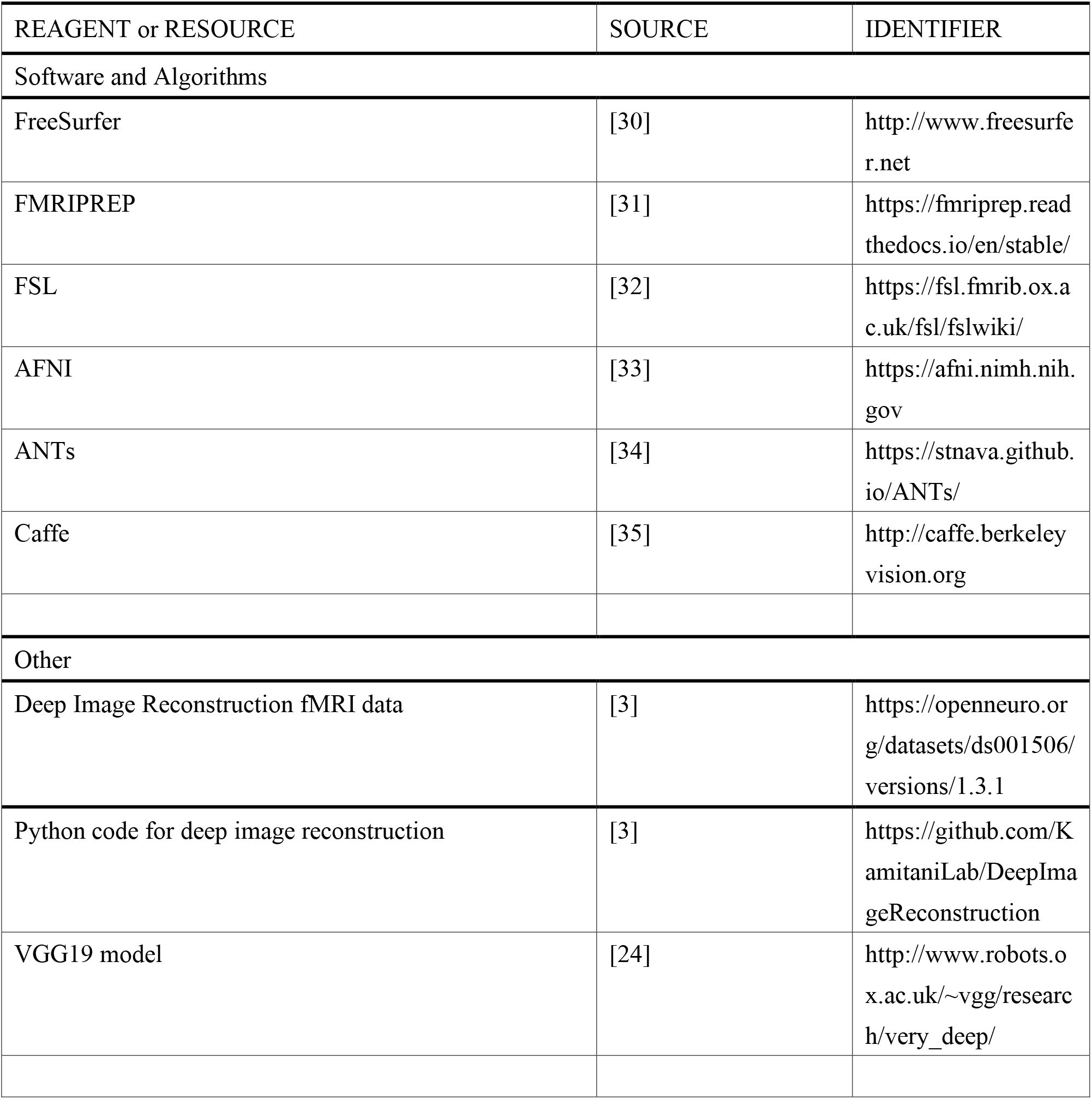

## CONTACT FOR REAGENT AND RESEROUCES SHARING

Further information and requests for resources and reagents should be directed to and will be fulfilled by the Lead Contact, Tomoyasu Horikawa.

## EXPERIMENTAL MODEL AND SUBJECT DETAILS

### Subjects

Five healthy subjects with normal or corrected-to-normal vision participated in our experiments: Subject 1 (male, age 34-36), Subject 2 (male, age 23-24), Subject 3 (female, age 23-24), Subject 4 (male, age 22-23), and Subject 5 (male, age 27-29). The first three subjects (Subjects 1-3) were the same with those in the previous study (Shen et al., 2019). For these subjects, we reused a subset of the previously published data (data for the training session, which was originally referred to as “training natural image session” of the “image presentation experiment”; available from https://openneuro.org/datasets/ds001506/versions/1.3.1), while newly collecting additional data (data for the test session). For the last two subjects (Subjects 4 and 5), we have newly collected a whole new dataset (data for the training session and the test session). The sample size was chosen based on previous fMRI studies with similar experimental designs [3, 22]. All subjects provided written informed consent for participation in the experiments, and the study protocol was approved by the Ethics Committee of ATR.

## METHOD DETAILS

### Stimuli

The stimuli consisted of color natural images, which were used in previous studies [3, 22] and were originally collected from the online image database *ImageNet* (2011, fall release) [36]. The images were cropped to the center and resized to 500 × 500 pixels.

### Experimental design

We conducted two types of experimental sessions: a training session and a test session. All stimuli were rear-projected onto a screen in the fMRI scanner bore using a luminance-calibrated liquid crystal display projector. The stimulus images were presented at the center of the display with a central fixation spot and were flashed at 2 Hz (12 × 12 and 0.3 × 0.3 degrees of visual angle for the visual images and fixation spot, respectively). To minimize head movements during fMRI scanning, subjects were required to fix their heads using a custom-molded bite-bar and/or a personalized headcase (https://caseforge.co/) individually made for each subject except for the case where subjects were reluctant to use those apparatuses (a subset of sessions with Subject 5). Data from each subject were collected over multiple scanning sessions spanning approximately 2 years. On each experimental day, one consecutive session was conducted for a maximum of 2 hours. Subjects were given adequate time for rest between runs (every 7-10 min) and were allowed to take a break or stop the experiment at any time.

### Training session

The training session consisted of 24 separate runs. Each run comprised 55 trials that consisted of 50 trials with different images and 5 randomly interspersed repetition trials where the same image as in the previous trial was presented (7 min 58 s for each run). Each trial was 8-s long with no rest period between trials. The color of the fixation spot changed from white to red for 0.5 s before each trial began, to indicate the onset of the trial. Additional 32- and 6-s rest periods were added to the beginning and end of each run, respectively. Subjects were requested to maintain steady fixation throughout each run and performed a one-back repetition detection task on the images, responding with a button press for each repeated image, to ensure they maintained their attention on the presented images. In one set of the training session, a total of 1,200 images were presented only once. This set was repeated five times (1,200 × 5 = 6,000 samples for training). The presentation order of the images was randomized across runs. This training session is identical to that conducted in the previous study [3] (referred to as “training natural image session” of the “image presentation experiment”). The data for the last two subjects (Subjects 4 and 5) were newly collected, whereas the data for the first three subjects (Subjects 1-3) were adopted from the data published by a previous study [3] (https://openneuro.org/datasets/ds001506/versions/1.3.1).

### Test session

The test session consisted of 16 separate runs. Each run comprised 55 trials that consisted of 10 single-image trials and 45 attention trials (7 min 58 s for each run). In each single-image trial, images were presented in the same manner as the training session. In each attention trial, subjects were presented with a sequence of images, each of which consisted of two successive cue images (2 s, 1 s for each cue) and spatially superimposed images of the two cue images (6 s), and were asked to attend to one image (indicated by green fixation shown with either of the two cue images) of a superposition of two images while ignoring the other such that the attended images are perceived more clearly. During the attention period, the subjects were also required to press one of two buttons gripped by their right hand to answer whether they correctly recognized which of the first and second cue image should be attended (percentages of correct, error, and miss trials among a total of 720 attention trials; 99.4%, 0.6%, and 0% for Subject 1; 98.8%, 0.6%, and 0.7% for Subject 2; 97.4%, 0.8%, and 1.8% for Subject 3; 99.9%, 0 %, and 0.1% for Subject 4; 93.5%, 3.5%, and 3.1% for Subject 5). In the test session, we used 10 out of 50 natural images that were used in the previous study [3] (“test natural image session” of the “image presentation experiment”; these images were not included in the stimuli of the training session). The 10 images were used to create a total of 45 combinations of superimposed images, and all these 45 unique superimposed images were presented as well as 10 unique single-images in each run with randomized orders (a total of 55 unique images were presented in each run). For each combination of superimposed two images, the number of trials to be the target of attention was balanced between the two images in every two consecutive runs and an entire session (a total of 8 trials for each).

### MRI acquisition

fMRI data were collected using a 3.0-Tesla Siemens MAGNETOM Verio scanner located at the Kokoro Research Center, Kyoto University. An interleaved T2*-weighted gradient-echo echo planar imaging (EPI) scan was performed to acquire functional images covering the entire brain (TR, 2000 ms; TE, 43 ms; flip angle, 80 deg; FOV, 192 × 192 mm; voxel size, 2 × 2 × 2 mm; slice gap, 0 mm; number of slices, 76; multiband factor, 4). T1-weighted (T1w) magnetization-prepared rapid acquisition gradient-echo (MP-RAGE) fine-structural images of the entire head were also acquired (TR, 2250 ms; TE, 3.06 ms; TI, 900 ms; flip angle, 9 deg; FOV, 256 × 256 mm; voxel size, 1.0 × 1.0 × 1.0 mm).

### MRI data preprocessing

We performed the MRI data preprocessing through the pipeline provided by FMRIPREP (version 1.2.1) [31]. For functional data of each run, first, a BOLD reference image was generated using a custom methodology of FMRIPREP. Using the generated BOLD reference, data were motion corrected using mcflirt from FSL (version 5.0.9) [32] and then slice time corrected using 3dTshift from AFNI (version 16.2.07) [29]. This was followed by co-registration to the corresponding T1w image using boundary-based registration implemented by bbregister from FreeSurfer (version 6.0.1) [30]. The coregistered BOLD time-series were then resampled onto their original space (2 × 2 × 2 mm voxels) using antsApplyTrainsforms from ANTs (version 2.1.0) [34] using Lanczos interpolation.

Using the preprocessed BOLD signals, data samples were created by first regressing out nuisance parameters from each voxel amplitude for each run, including a constant baseline, a linear trend, and temporal components proportional to the six motion parameters calculated during the motion correction procedure (three rotations and three translations). The data samples were temporally shifted by 4 s (2 volumes) to compensate for hemodynamic delays, were despiked to reduce extreme values (beyond ± 3 SD for each run), and were then averaged within each 8-s trial (training session, four volumes), last 6-s period of each trial (single-image trials in the test session, three volumes corresponding to second to fourth volumes in each trial), or 6-s attention period (attention trials in the test session, three volumes). For data from the test session, we discarded samples corresponding to error trials (miss or wrong button responses) from the main analyses unless otherwise stated (e.g., Figure S2C and D; numbers of samples after the removal, 716, 711, 701, 719, and 673 for Subject 1-5, respectively).

### Regions of interest (ROI)

V1, V2, V3, and V4 were delineated following the standard retinotopy experiment [37, 38]. The lateral occipital complex (LOC), fusiform face area (FFA), and parahippocampal place area (PPA) were identified using conventional functional localizers [39-41]. A contiguous region covering the LOC, FFA, and PPA was manually delineated on the flattened cortical surfaces, and the region was defined as the higher visual cortex (HVC). Voxels overlapping with V1–V3 were excluded from the HVC. Voxels from V1–V4 and the HVC were combined to define the visual cortex (VC).

### Deep neural network features

We used the Caffe implementation [35] of the VGG19 deep neural network (DNN) model [24], which was pre-trained with images in ImageNet [36] to classify 1,000 object categories (the pre-trained model is available from https://github.com/BVLC/caffe/wiki/Model-Zoo). The VGG19 model consisted of a total of sixteen convolutional layers and three fully connected layers. To compute outputs by the VGG19 model, all visual images were resized to 224 × 224 pixels and provided to the model. The outputs from the units in each of the 19 layers (immediately after convolutional or fully connected layers, before rectification) were treated as a vector in the following decoding and reconstruction analysis. The number of units in each of the 19 layers is the following: conv1_1 and conv1_2, 3211264; conv2_1 and conv2_2, 1605632; conv3_1, conv3_2, conv3_3, and conv3_4, 802816; conv4_1, conv4_2, conv4_3, and conv4_4, 401408; conv5_1, conv5_2, conv5_3, and conv5_4, 100352; fc6 and fc7, 4096; and fc8, 1000.

### Feature decoding analysis

We used a set of linear regression models to construct multivoxel decoders to decode a DNN feature pattern for a single presented image from a pattern of fMRI voxel values obtained in the training session (training dataset; samples from 6000 trials for each subject). The training dataset was used to train decoders to predict the values of individual units in feature patterns of all DNN layers (one decoder for one DNN unit). Decoders were trained using fMRI patterns in an entire visual cortex (VC) or individual visual subareas (V1–V4 and HVC), and voxels whose signal amplitudes showed the highest absolute correlation coefficients with feature values of a target DNN unit in the training data were provided to a decoder as inputs (with a maximum of 500 voxels).

The trained decoders were then applied to the fMRI data obtained in the test session (test dataset) to decode feature values of individual DNN units from fMRI samples constructed for each trial (samples from 160 single-image trials and 720 attention trials for each subject). Performances of the feature decoding were evaluated by calculating Pearson correlation coefficients between patterns of true and decoded feature values for each sample. To eliminate potential biases for calculating correlations due to baseline differences across units, feature values of individual units underwent z-score normalization using means and standard deviations of feature values of individual units estimated from the training data before calculating the correlations.

While similar decoding analyses were performed using a sparse linear regression algorithm in the previous studies [3, 22], we here used the least square linear regression algorithm as the number of training samples (6000 samples) exceeded the input dimensions (500 voxels). We have confirmed that results obtained from these algorithms were almost equivalent in decoding performances.

For the subsequent image reconstruction analysis, in order to compensate for possible differences in the distributions of true and decoded DNN feature values, the decoded feature values were normalized such that variances across units within individual channels/layers (groups of units within each channel for convolutional layers and all units within each layer for fully-connected layers) matched with the mean-variance of DNN feature values computed from independent 10,000 natural images. The feature values after this correction were then used as inputs to the reconstruction algorithm.

### Visual image reconstruction analysis

We performed the image reconstruction analysis using the previously proposed method [3], which optimizes pixel values of an input image based on a set of target DNN features such that the DNN features computed from the input image become closer to the target DNN features. The algorithm was originally formalized to solve the optimization problem for reconstructing images from image feature representations, such as activations of DNN units in a specific layer, by inverting them to pixel values for a certain reference image [42]. Shen et al. (2019) [3] extended the algorithm to combine features from multiple DNN layers and to use DNN features decoded from the brain instead of those computed from a reference image. To produce natural-looking images, they further introduced a deep generator network (DGN) [25], which was pre-trained to generate natural images using the generative adversarial network (GAN) framework [43], and performed optimization at the input space of the DGN.

In this study, following the method developed by the previous study [3], we used decoded DNN features from multiple DNN layers (a total of 19 layers of the VGG19 model) and introduced the pre-trained DGN [44] (the model for fc7 available from https://github.com/dosovits/caffe-fr-chairs) to constrain reconstructed images to have natural image-like appearances. The optimization was performed using a gradient descent with momentum algorithm [45] starting from zero-value vectors as the initial state in the latent space of the DGN (200 iterations; see [3] for details; code is available from https://github.com/KamitaniLab/DeepImageReconstruction).

### Evaluation of reconstruction quality

We evaluated the quality of reconstructed images by behavioral ratings to quantify the similarity of reconstructions to attended (for attention trials) or presented single images (for single-image trials) via a crowdsourcing platform. In this behavioral rating experiment, human raters were asked to judge which of a pair of two candidate images (an attended image and an unattended image for attention trials; a true [presented] image and a false image for single-image trials) is more similar to a reconstructed image. The evaluation was conducted by 20 raters for each reconstruction with a specific candidate pair (e.g., “post” and “leopard” for a reconstruction with target “post”), and a ratio of correct identification of attended and presented images among all raters (*n* = 20) and candidate pairs (*n* = 1 for attention trials, *n* = 9 for single-image trials) were defined as an accuracy of a reconstructed image. The evaluation was conducted for all reconstructed images from samples of attention and single-image trials. In Figure 2C and D, mean identification accuracies of attended and presented images averaged over all trials (*n* = 8 for attention trials, *n* = 16 for single-image trials) and paired-images (*n* = 2, attended/unattended for attention trials and true/false for single-image trials) are shown.

### Evaluation of visual appearance

To evaluate the visual appearance of stimulus images while paying attention to one of the overlapping images as in the fMRI test session (cf., Figure 1A; see Methods: “Experimental design”), we conducted an out-of-scanner behavioral experiment with an available subset of the subjects who participated in the fMRI experiments (Subject 1, 4, and 5). Each trial of this experiment consisted of a cue period (2 s), an attention period (6 s), a white-noise period (0.1 s), and an evaluation period (no time constraint), in which the cue and attention periods were the same as those in an attention trial in the fMRI test session. During a white-noise period, we presented white noise images (0.1 s, 60 Hz) on the same location of the presented images during the preceding cue and attention periods to diminish any potential effects of afterimages. During an evaluation period, we presented a test image consisted of a mixture of preceding two cue images, which was initialized with a random contrast for the weighted superpositions. Subjects were required to change the stimulus contrast of the presented test image to be closer to the visual appearance of the image perceived during the preceding attention period by pressing buttons for control. After matching the contrast, subjects were allowed to start the next trial in 2 s after pressing another button for proceeding. The evaluation was performed for all 45 combinations of superimposed images and two attention conditions (a total of 90 conditions), which were separately evaluated in two separate runs with randomized orders (~15 mins for each run). Each subject evaluated all conditions twice, and a mean contrast averaged across all subjects, repetitions, and attention conditions were used as a score for a specific pair (e.g., “owl” and “post”).

### Identification analysis

In the identification analysis based on feature correlations, correlation coefficients were calculated between a pattern of decoded features and patterns of image features calculated from two candidate images (one for attended and the other for unattended images for attention trials; one for true [presented] and the other for false images for single-image trials). For each reconstructed image from attention trials, the pair-wise identification was performed with a pair of attended and unattended images (one pair for each sample). For each reconstructed image from single-image trials, the identification was performed for all pairs between one true (presented) image and the other nine false images that were used in the test session (nine pairs for each sample). The image with a higher correlation coefficient was selected as the predicted image. The accuracy of each sample was defined by the proportions of correct identification. To eliminate potential biases due to baseline differences across units, feature values of individual units underwent z-score normalization using means and standard deviations of feature values of individual units estimated from the training data before performing the identification.

## QUANTIFICATION AND STATITICAL ANALYSIS

One-sided Wilcoxon signed-rank test was used to test the significance of the identification accuracies based on behavioral evaluations (*n* = 90; Figure 2C), and to test the significance of the single-image identification accuracies based on decoded DNN features (*n* = 160; Figure 4B right). A correlation between the identification accuracies of attention and single-image trials evaluated based on behavioral evaluations was tested by *t*-test (*n* = 45; Figure 2D). One-sided binomial test was used to test the significance of the attended image identification accuracies based on decoded DNN features (*n* = 716, 711, 701, 719, and 673 for Subject 1-5, respectively; Figure 4B left).

## DATA AND SOFTWARE AVAILABILITY

The experimental code and data that support the findings of this study are available from our repository (code for feature decoding: https://github.com/KamitaniLab/GenericObjectDecoding, code for image reconstruction: https://github.com/KamitaniLab/DeepImageReconstruction), and open data repository (OpenNeuro: https://openneuro.org/datasets/ds003430).

## Supplemental Information

### Supplemental Figures

Figure S1. Feature decoding and reconstruction results for single-image trials. Related to Figure 2.

Figure S2. Examples of reconstructed images for attention trials. Related to Figure 2.

Figure S3. Correlation coefficients between decoded feature patterns and image feature patterns computed from weighted superpositions for individual subjects. Related to Figure 3.

Figure S4. Identification accuracy from attention and single-image trials. Related to Figure 4.

### Supplemental Video

Video S1. Reconstructions of attended images. Related to Figure 2.

**Figure S1.**
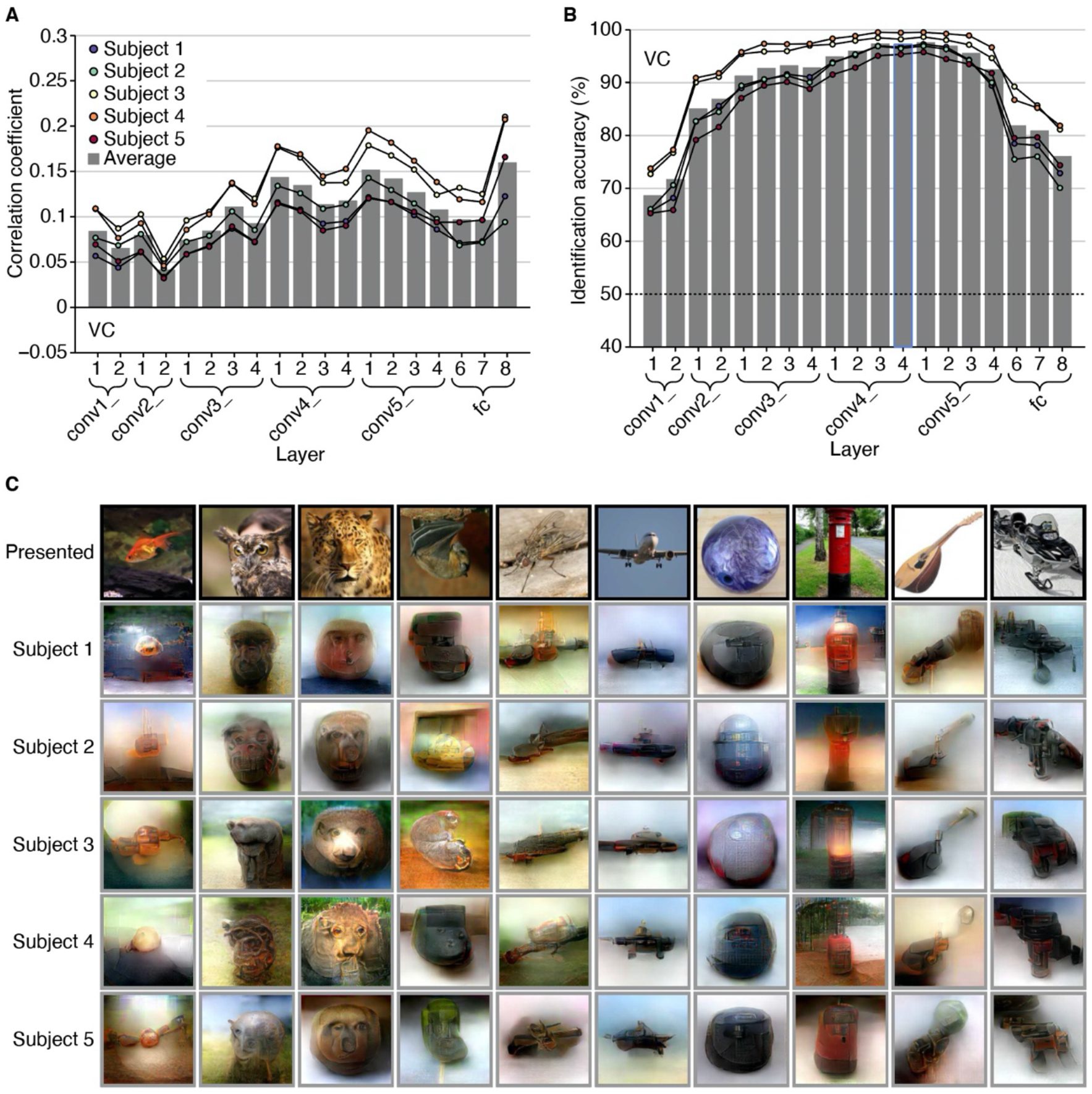
Feature decoding and reconstruction results for single-image trials. Related to Figure 2. (A) Feature decoding accuracy for single-image trials. A decoding accuracy was evaluated by calculating a Pearson correlation coefficient between a pattern of decoded feature values and a pattern of image feature values computed from presented images for each sample (decoded from the visual cortex [VC]). Correlation coefficients were averaged across samples from the single-image trials (a total of 160 trials for each subject, colored dots), and the mean correlations averaged across subjects (gray bars) are shown for each layer of the VGG19 model. (B) Pair-wise image identification accuracy for single-image trials. Identification accuracy obtained by the pair-wise identification analysis is shown for each layer of the VGG19 model (decoded from VC; chance level, 50%; see Methods: “Identification analysis”). In the analysis, correlation coefficients were calculated between a pattern of decoded features and patterns of image features of two candidate images (one for true [presented], and the other for false), and the image with a higher correlation coefficient was selected as the predicted image. For each sample, the pair-wise identification was performed for all pairs between one true image and the other nine false images used in the test session (nine pairs for each sample). The accuracy of each sample was defined by the proportions of correct identification. Conventions are the same with (A). (C) Examples of reconstructed images from single-image trials. The reconstructed images produced from samples of each of the single-image trials are shown for five subjects (decoded from VC). Conventions are the same with Figure 2B. These high feature decoding accuracy and reconstruction quality for samples of single-image trials validated the model performances for all subjects.

**Figure S2.**
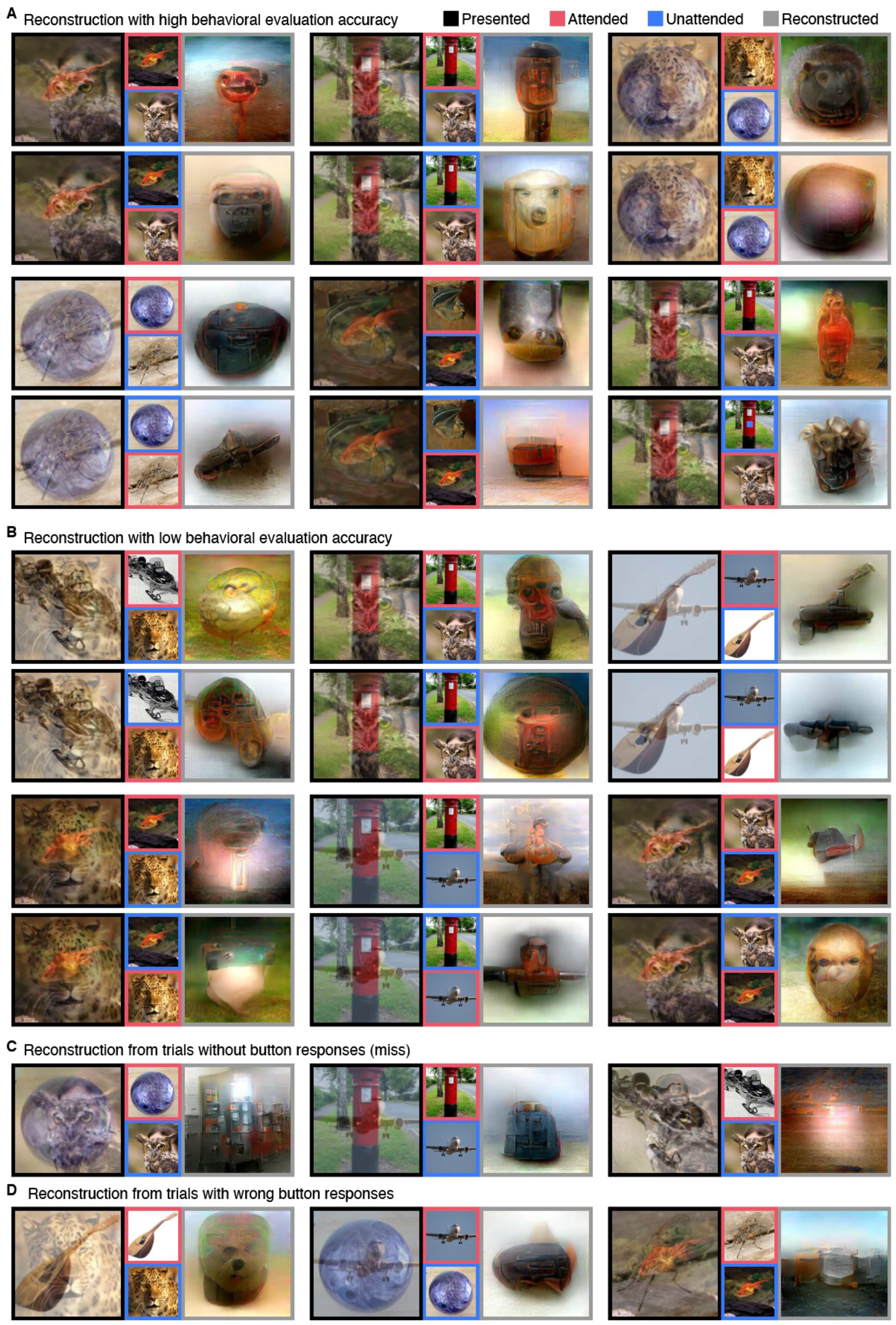
Examples of reconstructed images for attention trials. Related to Figure 2. (A) Examples of attended image reconstructions with high behavioral rating accuracies. Reconstructed images with relatively high rating accuracies (higher than 80%) are shown. Conventions are the same with Figure 2A. (B) Examples of attended image reconstructions with low rating accuracies. Reconstructed images with relatively low rating accuracies (lower than 60%) are shown. The failures of attended image reconstructions were categorized into clutter images, mixtures of two superimposed images, or images more similar to unattended images. (C) Reconstructed images from samples for trials without button responses. Reconstructed images obtained from samples for miss trials, in which subjects missed to press a button to indicate correct recognition of target images, are shown. The reconstructions from these miss trials tended to be not similar to either of the two superimposed images. (D) Reconstructed images from samples for trials with wrong button responses. Reconstructed images obtained from samples for error trials, in which subjects incorrectly pressed a button to indicate target images, are shown. The reconstructions from these error trials sometimes produced images judged to be similar to non-target (or instructed to be unattended) images.

**Figure S3.**
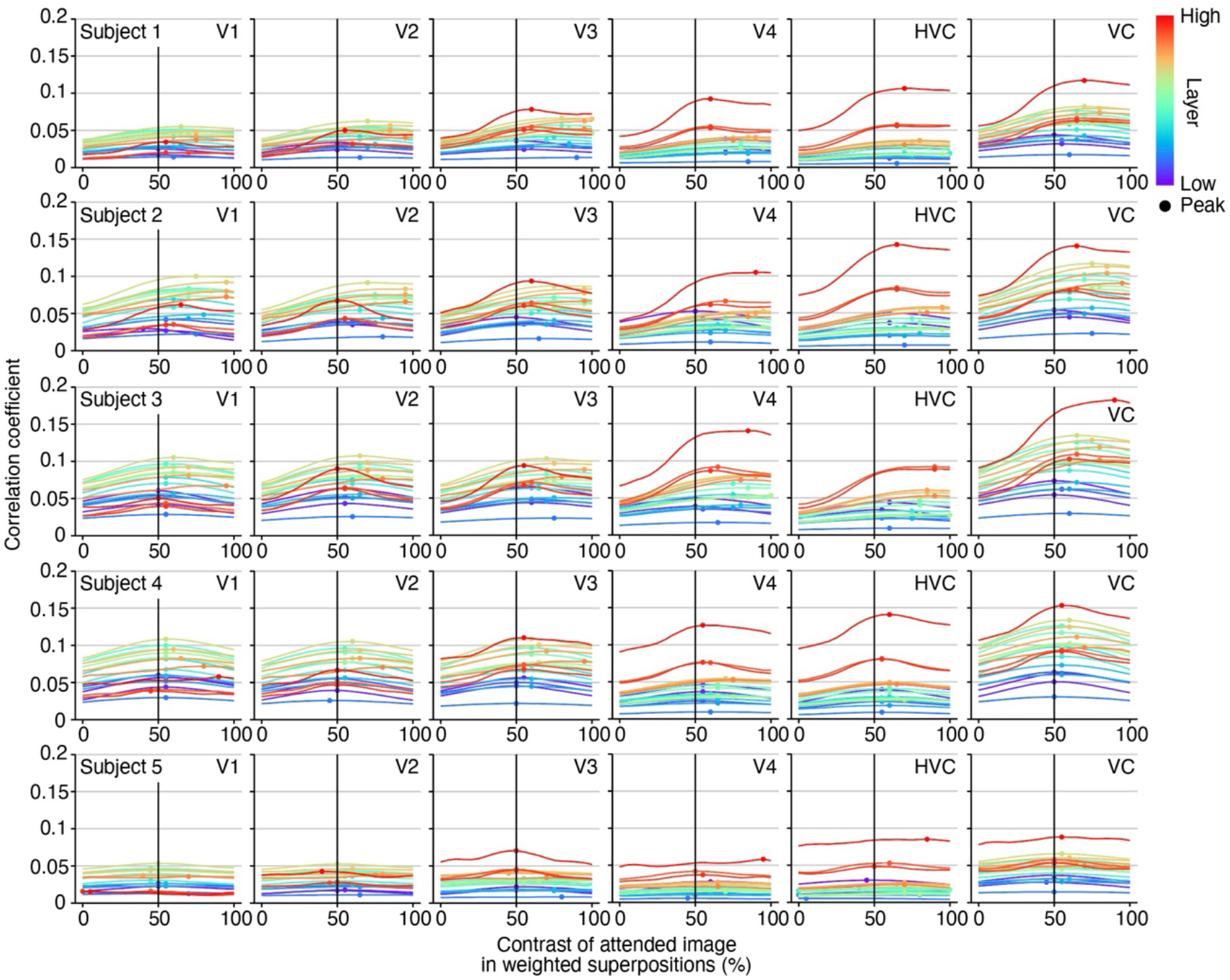
Correlation coefficients between decoded feature patterns and image feature patterns computed from weighted superpositions for individual subjects. Related to Figure 3. Correlations between decoded feature patterns and image feature patterns computed from weighted superpositions with different contrasts are shown for individual subjects. Conventions are the same with Figure 3C. The subjects whose reconstructions from attention trials were highly evaluated (e.g., Subject 1-3; cf., Figure 2C) showed greater biases in decoded feature patterns, which deviated toward attended images (100%) from presented images (50%).

**Figure S4.**
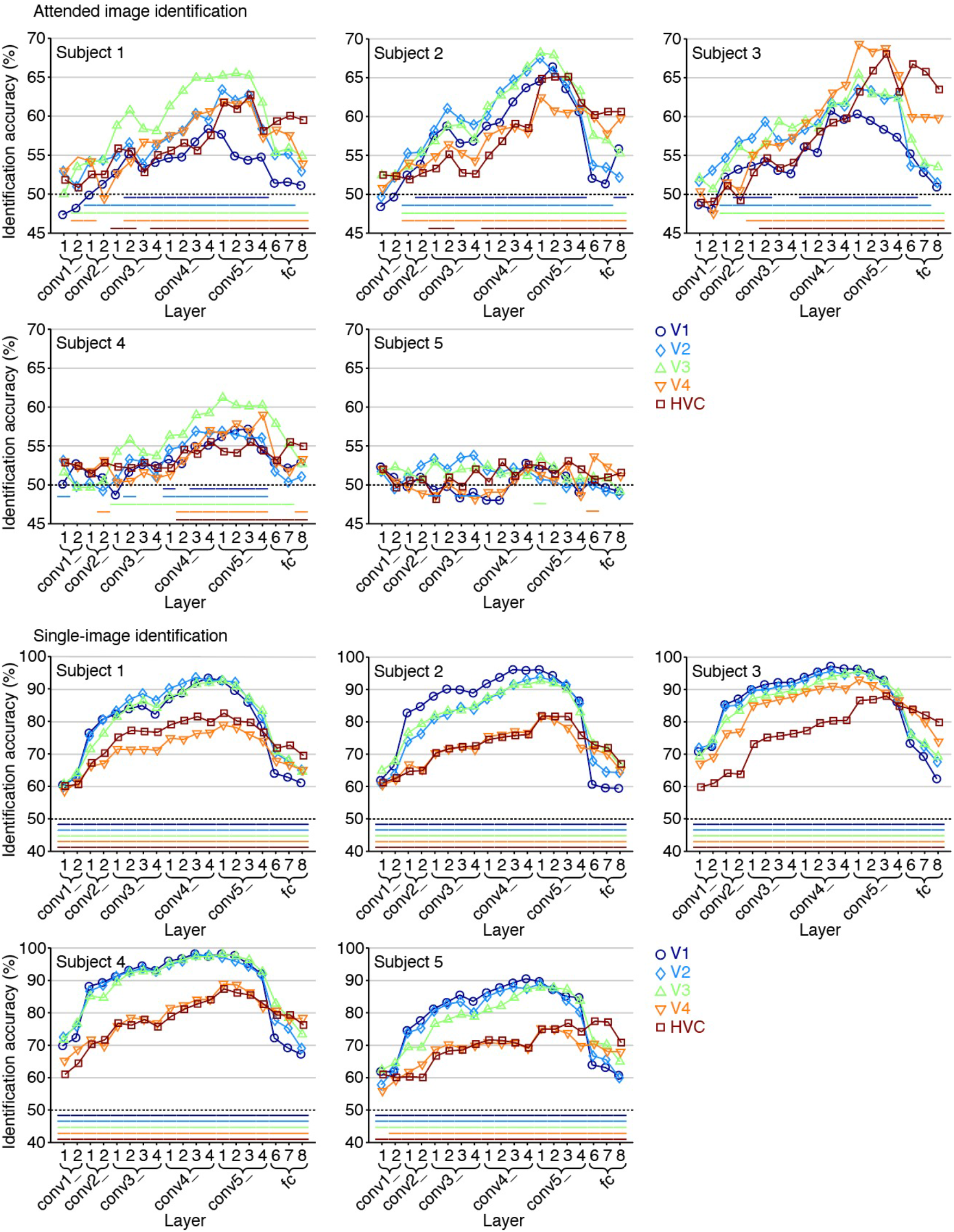
Identification accuracy from attention and single-image trials. Related to Figure 4. Colored lines beneath data indicate results of statistical tests (for attention, one-sided binomial test, *p* < 0.05; for single-image, one-sided Wilcoxon signed-rank test, *p* < 0.05). The results showed relatively larger variabilities among subjects in the accuracies of the attended image identification than those of the single-image identification, possibly due to the individual differences of the ability to direct their selective attention. Differences of brain regions that showed high attended image identification accuracies might be attributable to the differences of their strategies for attention, as we did not explicitly provide specific strategies for their attempt of attention.

**Video S1. Reconstructions of attended images. Related to Figure 2**. The iterative optimization process is shown for reconstructions from attention trials (the last 80 steps of a total of 200 optimization steps; left, presented image; center, attended or unattended image; red frame, attended image; right, reconstructed image; cf., Figure 2; https://youtu.be/iJAF8d7d9dc).

